# Future carbon emissions from global mangrove forest loss

**DOI:** 10.1101/2020.08.27.271189

**Authors:** M.F. Adame, R.M. Connolly, M.P. Turschwell, C.E. Lovelock, L. Fatoyinbo, D. Lagomasino, L.A. Goldberg, J. Holdorf, D.A. Friess, SD. Sasmito, J. Sanderman, M. Sievers, C. Buelow, B.J. Kauffman, D. Bryan-Brown, C.J. Brown

## Abstract

Mangroves have among the highest carbon densities of any tropical forest. These “blue carbon” ecosystems can store large amounts of carbon for long periods, and their protection reduces greenhouse gas emissions and supports climate change mitigation. The incorporation of mangroves into Nationally Determined Contributions to the Paris Agreement and their valuation on carbon markets requires predicting how the management of different land-uses can prevent future greenhouse gas emissions and increase CO_2_ sequestration. Management actions can reduce CO_2_ emissions and enhance sequestration, but should be guided by predictions of future emissions, not just carbon storage. We project emissions and forgone soil carbon sequestration potential caused by mangrove loss with comprehensive global datasets for carbon stocks, mangrove distribution, deforestation rates, and drivers of land-use change. Emissions from mangrove loss could reach 2,397 Tg CO_2eq_ by the end of the century, or 3,401 Tg CO_2eq_ when considering forgone carbon sequestration. The highest emissions were predicted in southeast and south Asia (West Coral Triangle, Sunda Shelf, and the Bay of Bengal) due to conversion to aquaculture or agriculture, followed by the Caribbean (Tropical Northwest Atlantic) due to clearing and erosion, and the Andaman coast (West Myanmar) and north Brazil due to erosion. Together, these six regions accounted for 90% of the total potential CO_2eq_ future emissions. We highlight hotspots for future emissions and the land-use specfic management actions that could avoid them with appropriate policies and regulation.

## Introduction

The capacity of mangroves to store carbon and mitigate greenhouse gas emissions became prominent a decade ago (Donato et al., 2011). Since then, mangroves have gained international interest for their potential to contribute to carbon mitigation strategies and for their ecosystem services that support adaptation to climate change (Lovelock & Duarte, 2019). Hundreds of site-scale studies have been conducted to understand the distribution and accumulation of mangrove soil carbon and aboveground biomass (Kauffman et al., 2020). These site-scale models have supported globally comprehensive spatial models of carbon storage (Rovai et al., 2018; Sanderman et al., 2018; Simard et al., 2019). Simultaneously, global efforts to accurately map and monitor mangrove cover and health have provided unprecedented knowledge on the risks that mangrove forests face (Bunting et al., 2018; Goldberg et al., 2020; Hamilton & Casey, 2016). These studies have enabled global-scale estimation of mangrove carbon storage and its historical loss across different nations (Murdiyarso et al., 2015; Serrano et al., 2019) and globally (Atwood et al., 2017).

Improved management can reduce CO_2_ emissions from mangrove forest loss and enhance the sequestration potential of disturbed forests (Friess et al., 2020; O’Connor et al., 2020), but management actions should be guided by predictions of future emissions, not just carbon storage. The effectiveness of management relies on understanding how much emissions can be avoided by specific actions, for instance, by reducing land conversion or by increasing restoration efforts. Predictions of CO_2_ emissions from mangrove loss linked with specific land-use changes can underpin the selection of actions to support adequate mangrove management actions for specific drivers of loss. These actions include improving the representation of mangroves in the National Determined Contributions (NDCs) committed to in the Paris Climate Agreement, strengthening their role as natural-based solutions, and improving their valuation on carbon markets (Adame et al., 2018; Seddon et al., 2019).

Recent advances in mapping of mangrove area, rates of loss, carbon storage and emission factors now enable predictions of CO_2_ emissions at the global scale (Worthington et al., 2020). These predictions should overcome several key limitations in past studies (Macreadie et al., 2019). First, carbon emission estimates have yet to associate particular land-use changes with CO_2_ emissions, as mapping of global drivers of mangrove loss has just recently become available (Goldberg et al., 2020). Second, many global estimates have included only the first meter of soil, thus, underestimating the total carbon content of mangroves and the emissions that arise from their conversion to other land uses (Kauffman et al., 2020). Third, estimates of global carbon emissions have not included the forgone carbon sequestration and they do not account for the lost opportunity of sequestration when mangroves are lost (Maxwell et al., 2019). And finally, global estimates have treated all CO_2_ emissions from mangroves as occurring in the year of loss (Atwood et al., 2017). Depending on the type of land-use change and the carbon pool affected, it can take years or even decades for the carbon stored in mangroves to be emitted into the atmosphere (Lovelock et al., 2017) and exported through tidal exchange (Maher et al., 2013).

To overcome current limitations in global estimations, we developed a spatial model that projects emissions caused by mangrove loss. Our model synthesised information from multiple newly available global datasets, including carbon stocks (Kauffman et al., 2020; Sanderman et al., 2018; Simard et al., 2019), mangrove distribution (Bunting et al., 2018), deforestation rates (Hamilton & Casey, 2016), drivers of land-use change (Goldberg et al., 2020) and emissions factors (Sasmito et al., 2019). We provide predictions of future CO_2_ emissions from mangrove loss, accounting for the effect of proximate drivers of land-use change including: a) conversion to commodities, such as agriculture or aquaculture, b) coastal erosion, c) clearing, d) extreme climatic events, and e) conversion to human settlements (Goldberg et al., 2020). Importantly, we account for the foregone opportunity of soil carbon sequestration when mangroves are lost (Maxwell et al., 2019). Our modelled emissions reflect the realistic temporal scale of emissions, which are not annual, but decadal (Lovelock, et al., 2017). By linking emissions with specific land-use changes and accounting for future removals, we provide for the first time, spatially explicit information on how different drivers of mangrove loss are causing CO_2_ emissions and what management actions can prevent them.

## Methods

### Mangrove area, rates of loss and drivers of change

We divided global mangrove extent (Bunting et al., 2018) into the marine provinces (top-level category of the bioregions) that contained mangroves (Spalding et al., 2007; Van der Stocken et al., 2019; Fig. 1, S1, Table S1). We selected this approach to estimate global CO_2_ emissions because it is well aligned with climatic and geomorphic characteristics of mangroves, which are variables associated with carbon stocks and losses (Dürr et al., 2011; Rogers et al., 2019). Deforestation rate for each province was obtained from the dataset by Hamilton and Casey (2016) for the years 2000-2012.

**Figure. 1.**
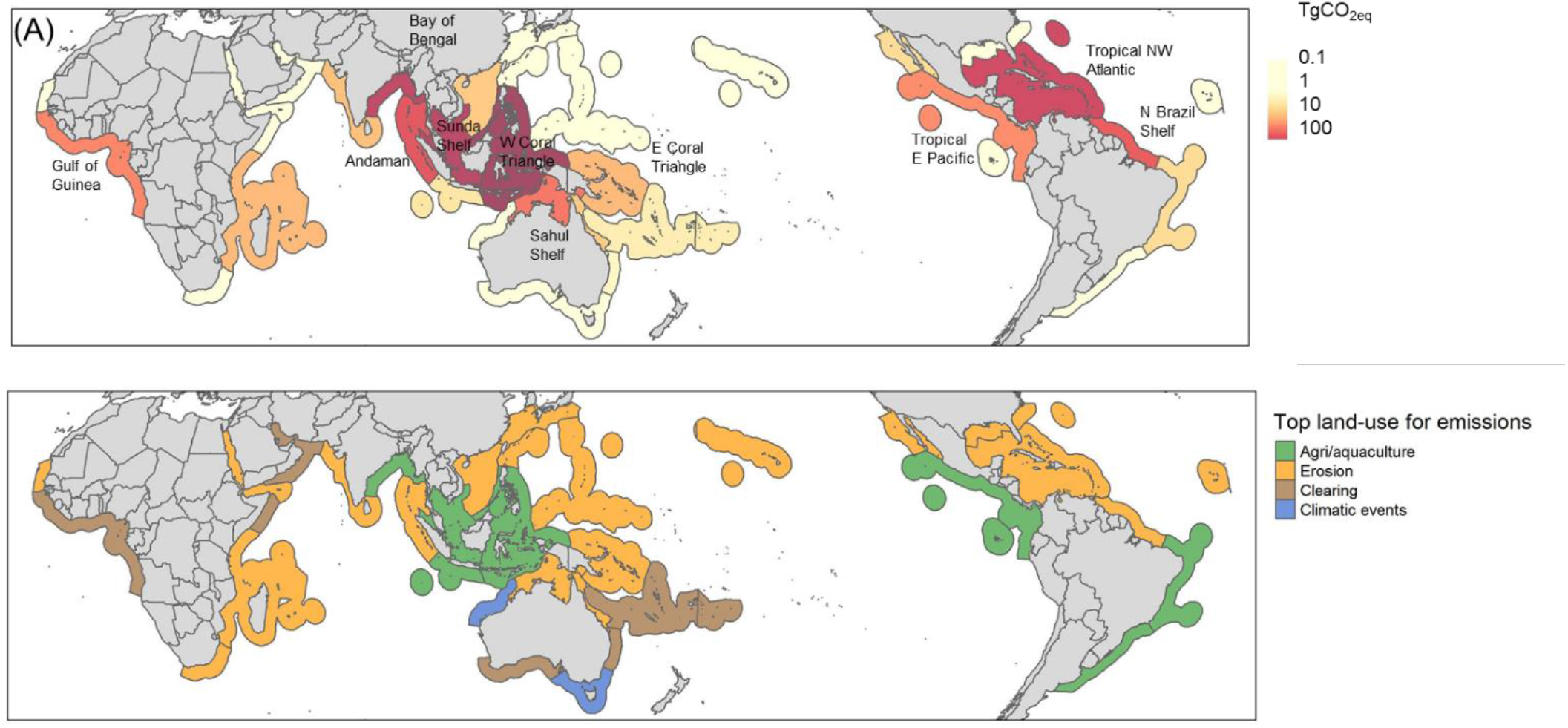
(A) Global projected CO_2 eq_ emissions (Tg) by the end of the century (2010-2100) for the marine provinces of the world, and (B) the proximate driver responsible for the largest CO_2_ emissions for each marine province (Goldberg et al. 2020). The names for all marine provinces can be found in Fig. S1 and Table S1.

The drivers of mangrove loss for each province (2000-2016) were obtained from changes in mangrove area and a decision-tree model that separated drivers into five categories: a) conversion to commodities, such as agriculture or aquaculture, b) coastal erosion, c) clearing due to various activities including logging or hydrological modifications, d) extreme climatic events, such as tropical storms and fluctuations in sea level, and e) conversion to human settlements (Goldberg et al., 2020). Briefly, mangrove loss was estimated from the Surface Reflectance Tier-1 Landsat 5TM, 7ETM+, and 8OLI imagery within Google Earth Engine. A baseline period (1999-2001) of a Normalised Difference Vegetation Index (NDVI) optimised mosaic representing the year 2000 was created from where mangrove area change was estimated. A threshold change value of −0.2 that occurred within the Mangrove Forests of the World extent (Giri et al., 2011) was used to indicate the areas of mangroves that had transitioned from forest to no-forest (Lagomasino et al., 2019). A random forest classification was applied to the areas showing a drop in NDVI greater than or equal to 0.2. These areas were trained for each land cover type: water, dark soils, and bright soils. Erosion was defined as a transition to water that intersected rivers and coastlines. Commodities (agriculture/aquaculture) were defined where mangrove loss intersected the Global Food Security-support Analysis Data Cropland Extent 30-m (GFSAD-30) layer (www.usgs.gov/centers/wgsc/science/global-food-security-support-analysis-data-30-m). Human settlements were defined as the bright soil land cover class that intersected with the Global Human Settlements Layer (GHSL). Clearing or non-productive conversions were defined at bright and dark soil land cover that intersected with a 5 km buffer around the GRIP-4 global roads dataset (//doi.org/10.7927/H4VD6WCT) and the GHSL human settlement dataset (//ghsl.jrc.ec.europa.eu/data.php). Lastly, conversion by extreme climatic events was defined as all other areas of mangrove loss that did not occur within a 5 km infrastructure buffer.

### Total Ecosystem Carbon Stocks (TECS)

Soil organic carbon (SOC) stocks for one and two meters of soil were obtained from the global SOC dataset (Sanderman et al., 2018), which was derived from a random forest model trained on field measurements. For aboveground ecosystem carbon (ABC), biomass was obtained from the global dataset of mangrove biomass (Simard et al., 2019). The total biomass per province was divided by mangrove area to obtain a mean ABC per province and multiplied by a factor of 0.48 to obtain carbon values (Kauffman & Donato, 2012). We compared the TECS obtained from the global models with field measurements from provinces where data was available (Kauffman et al., 2020) with a linear regression (IBM SPSS Statistics, v25). TECS obtained from global models (Sanderman et al., 2018; Simard et al., 2019) were lower in provinces with high stocks (> 1,200 MgC ha^−1^, e.g. Sunda Shelf and West Coral Triangle) and higher in provinces with small stocks (< 220 MgC ha^−1^, e.g. Northwest Australian Shelf and Somali Arabian). There were only 15 provinces with field data, but the model predictions for these provinces were very close to the modelled data when the depth of SOC was selected at two meters (Fig. S2, Table S1). Hence, we calculated TECS for all provinces as the sum of ABC and SOC for the top two meters of soil.

### Emissions factors

The emission factor is the fraction of carbon that is emitted given conversion to a specific land-use. We selected an emission factor for each province and activity from a recent global systematic review (Sasmito et al., 2019). Each emission factor was given a level of confidence (Table S2) from low to high, with Level 1 (lowest confidence) given to emission factors obtained from a global average, specific to that proximate driver; Level 2 to those obtained from a similar region, specific to that proximate driver; and Level 3 (highest confidence), from a similar region with the same geomorphic setting, specific to that proximate driver (Dürr et al., 2011).

### Model for projecting emissions and missed opportunities to sequester carbon

We updated a model of carbon emissions from deforested mangroves (Adame et al., 2018) to account for drivers of land-uses and soil carbon sequestration. The model allowed for variable carbon stocks across discrete spatial units and assumed a constant rate of deforestation and a constant rate of emissions once mangroves were lost. We modelled forgone carbon sequestration from mangrove loss in each province as the difference between carbon storage with deforestation and a counterfactual with no deforestation:

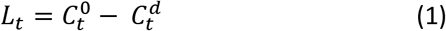

Cumulative carbon emissions, *L_t_*, were described by three dynamic equations:

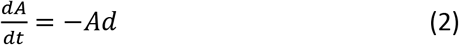

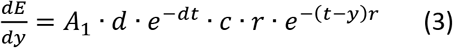

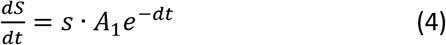

Where A is the area of mangroves in hectares, d is the total deforestation rate across all land-uses, E is the emissions, r is the rate of emissions from deforested mangroves, c is the total carbon stock emitted per hectare, y is the year of deforestation, S is sequestered carbon and s is the yearly sequestration rate per hectare.

We assumed that future rates of deforestation due to each of the five drivers were in proportion to their historical contributions to loss from Goldberg et al. (2020). Therefore, province-specific potential emissions per hectare were scaled by land-use types and their respective emission factors:

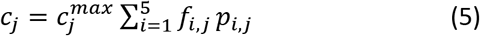

Where 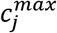 is maximum labile carbon per hectare for a province including SOC and AGC, *f_i,j_* are province and land-use specific emissions factors, and *p_i,j_* are the proportional contributions of each land-use type to past deforestation.

#### Sensitivity analyses to global datasets and model robustness

To determine how sensitive our future predictions were to each of the variables selected, we conducted sensitivity analyses where we repeated the predictions with different datasets for mangrove area (Bunting et al., 2018; Hamilton & Casey, 2016), sources of data (modelled and field) (Kauffman et al., 2020; Sanderman et al., 2018; Simard et al., 2019), and SOC depth (1m, 2m, and whole sediment column).

We also conducted sensitivity analyses on the emissions factors relating to erosion and extreme climatic events, which can be highly variable. Erosion can cause large emission in one location but partly compensate the mangrove area loss by accretion in other areas (Lagomasino et al., 2019). Extreme climatic events, such as tropical storms can cause large-scale mortality; however, some areas can naturally recover after a few years if conditions are appropriate, thereby reducing emissions (Krauss & Osland, 2020). Thus, we implemented the model with emission factors of 50 and 100% for erosion, and with and without the mangrove area loss from climatic events. Finally, we conducted further formal sensitivity analyses of the model to all the parameter inputs by taking the derivative of *L_t_* (cumulative carbon emissions) with respect to each parameter.

## Results and Discussion

### Inputs to the model

First, we present summaries of the input data, noting that this data has been reported elsewhere, but not aggregated by provinces. The mean TECS (mean ± SE, [range]) measured in the field for all provinces was 624.5 ± 96.9 Mg C ha^−1^ (181.5–1,434.9). The mean modelled SOC in the top meter of soil was 331.3 ± 74.9 (207.4–497.8) Mg C ha^−1^, in the top two meters was 646.7 ± 150.6 (408.6–975.9) Mg C ha^−1^, and mean ABC was 101.2 ± 93.5 (9.9–466.0) Mg C ha^−1^. The provinces with the largest area of mangroves were West Coral Triangle, the Gulf of Guinea, Sahul Shelf, and Tropical Northwest Atlantic (Table S1, Fig. S3). Ten provinces contained 88% of all the mangroves in the world. From 2000 to 2012, 35 of the 37 provinces had some level of deforestation, with mean annual losses of 0.09 ± 0.02%. The highest deforestation rates were in the Bay of Bengal (0.54%), Sunda Shelf (0.34 %), West Coral Triangle (0.32%), and Tropical Northwest Atlantic (0.14%) (Table S1, Fig. S3).

Conversion of mangroves to aquaculture/agriculture was the primary proximate driver of mangrove loss, which caused the conversion of 219,392 ha of mangroves from 2000 to 2016, especially in the West Coral Triangle (100,231 ha), Bay of Bengal (54,602 ha) and Sunda Shelf (44,530 ha, Fig. 1A). This corresponds to 87.1, 73.5, and 70.4% of their total mangrove loss, respectively. The second most important proximate driver of mangrove loss was erosion, which caused the loss of 92,787 ha, mainly in North Brazil Shelf (20,547 ha, 54.9% of the total mangrove loss), the Bay of Bengal (14,309 ha, 19.3%) and Sunda Shelf (11,863 ha, 18.8%). The third proximate driver of mangrove loss was extreme climatic events, causing the loss of 41,525 ha of mangroves, mainly in North Brazil Shelf (8,605 ha, 5.7%), Tropical Northwest Atlantic (8,257 ha, 30.7%) and Sahul Shelf (8,480 ha, 42.2%). The fourth most important driver was mangrove clearing which caused the loss of 39,595 ha, mostly in Tropical Northwest Atlantic (9,244 ha, 30.7%), Gulf of Guinea (8,738 ha, 42.2%), and West Indian Ocean (5,817 ha, 35.6%). Finally, the fifth proximate driver of mangrove loss was human settlement, which caused the loss of 10,529 ha, mostly in the Sunda (3,980 ha, 6.3%) and Sahul Shelf (3,397 ha, 16.4%).

### Predictions of carbon emissions and lost opportunities to sequester carbons

Global emissions from mangrove loss are projected to reach 2,397 TgCO_2 eq_ by the end of the century (2020-2100). Including the loss of potential carbon sequestration once mangroves are deforested (considered to have a global mean value of 1.5 MgC ha^−1^ yr^−1^; Alongi, 2014) increased our projection to 3,401 TgCO_2 eq_. Previous estimates of mangrove emissions for the same period varied enormously, between 630 and 40,230 TgCO_2eq_ (Friess, et al., 2020). Our projection lies towards the lower end of this range, and we consider it more accurate because of the inclusion of land-use drivers, time lags, and foregone future sequestration that were not considered in previous studies. Projected CO_2_ emissions showed significant geographical variability (Fig 1A). They were highest for the West Coral Triangle (712 TgCO_2 eq_), followed by Sunda Shelf (452 TgCO_2 eq_), Bay of Bengal (369 TgCO_2 eq_), Tropical Northwest Atlantic (312 TgCO_2 eq_), Andaman coast (161 TgCO_2 eq_), and North Brazil Shelf (137 TgCO_2 eq_). Collectively, these provinces contributed 90% of the total projected global CO_2_ emissions (Fig. 1A, Fig. 2, Table S3).

**Figure 2.**
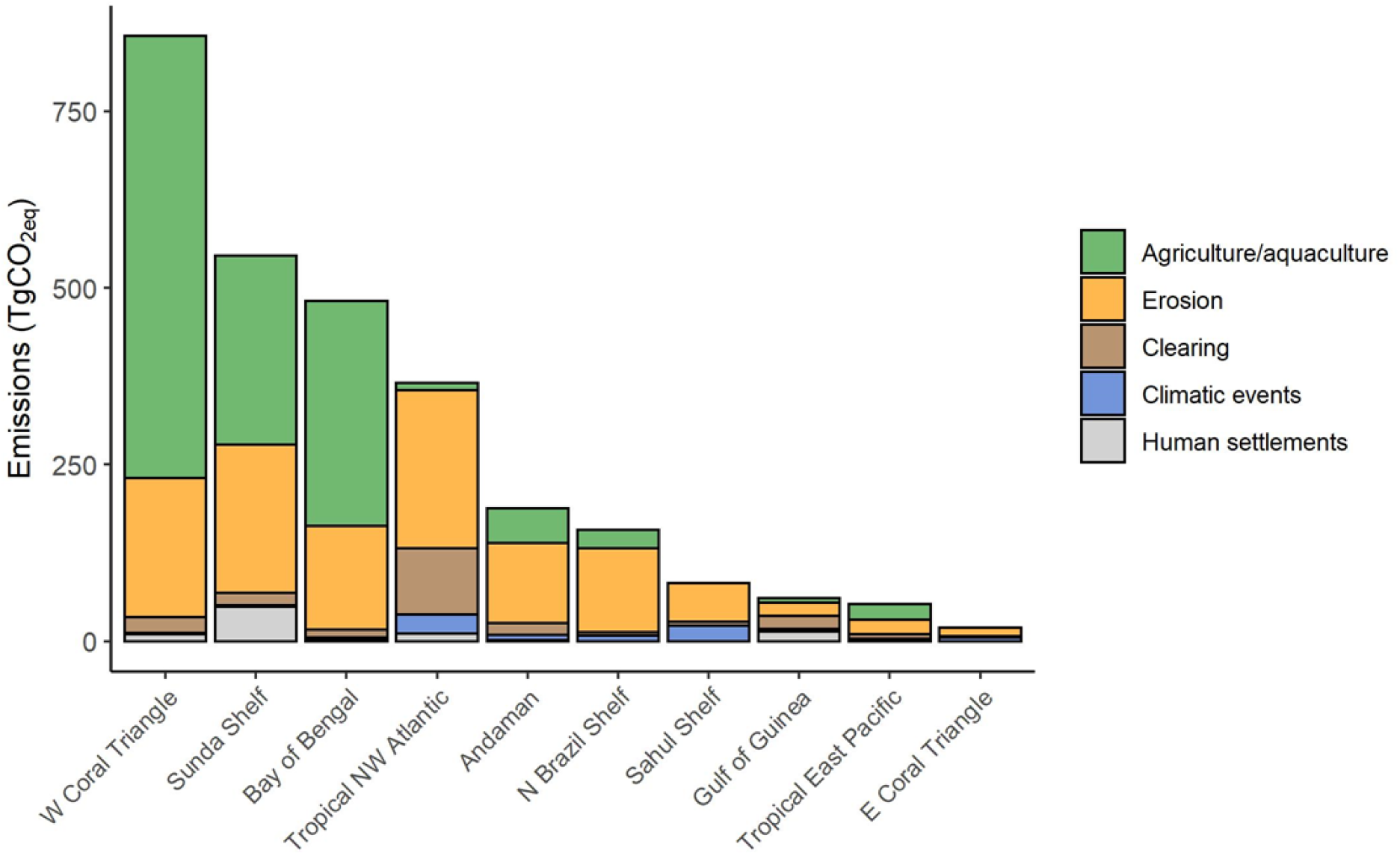
Cumulative CO_2 eq_ emissions (Tg) by the end of the century (2010-2100) attributed to the proximate drivers of mangrove loss for the marine provinces ranked in the top ten for future CO_2_ emissions.

The West Coral Triangle, Sunda Shelf, and the Bay of Bengal had the highest predicted emissions due to mangrove conversion to agriculture/aquaculture at 985 Tg CO_2 eq_ (Fig. 1B, 2). This region has been previously highlighted as a global hotspot of mangrove CO_2_ emissions (Atwood et al., 2017). Within these provinces, clearing of large areas of carbon-rich mangroves has occurred for rice, oil palm, aquaculture, and rubber plantations (De Alban et al., 2019; Richards & Friess, 2016). In Indonesia, the conversion of mangroves to aquaculture contributed almost 15% of their national emissions (Murdiyarso et al., 2015). In Myanmar, deforestation of mangroves has been driven by national policies that support the intensification of rice production to increase food security (Webb et al., 2014). Our predictions suggest that emissions from the south and southeast East Asia will be the highest globally by the end of the century due to the intensity of land-use changes and large mangrove carbon stocks (Fig. S3). These emissions have the potential to be managed through changes in agricultural practices, and the restoration of formerly converted mangrove areas, such as disused aquaculture ponds and land with saltwater intrusion. For instance, the improvement of the agriculture/aquaculture industry in the West Coral Triangle could reduce up to 73% of the projected CO_2_ emission for this region (Fig. 3A).

**Figure 3.**
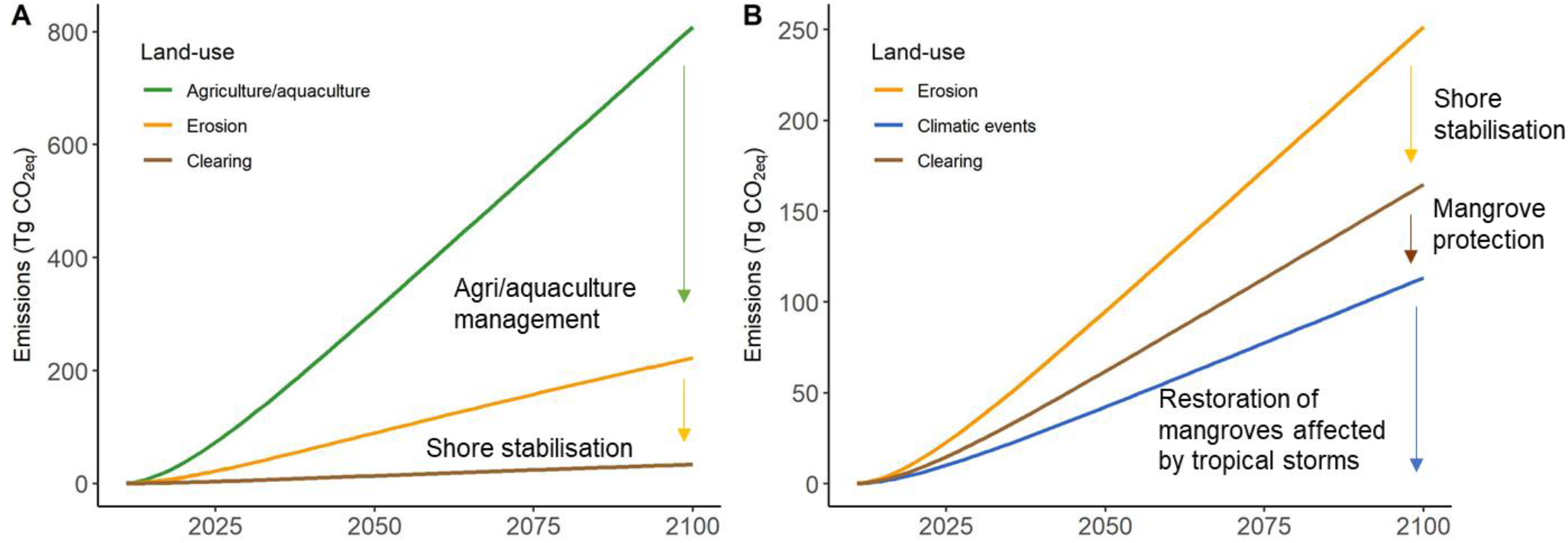
Emission reductions (Tg CO_2 eq_) that could be achieved from (A) management of agriculture/aquaculture and shore stabilisation in the West Coral Triangle, and (B) decrease in erosion through shore stabilisation, mangrove protection to avoid clearing, and management of mangroves affected by tropical storms in the Tropical Northwest Atlantic.

An important driver of mangrove loss in the southeast and southeast Asia was erosion, which accounted for 23, 38, and 30% of the total emissions from the West Coral Triangle, Sunda Shelf, and Bay of Bengal provinces, respectively (Fig. 2). Additionally, the adjacent province of Andaman (west Myanmar, Bangladesh, and east India) had significant emissions due to erosion (98 TgCO_2 eq_ or 60% of its total emissions). In the Sundarbans, changes in river flows have reduced sediment inputs, which caused the loss of over 7,500 ha of coastline in the last 37 years (Bhargava et al., 2020). In areas prone to high rates of erosion, decreasing emissions would need to be achieved through shore stabilisation and the management of rivers and dams to provide sediment inputs that support the maintenance of mangrove surface elevations and habitat area (Lovelock et al., 2015; Fig. 3A). Landward migration of mangroves, if coastal squeeze is avoided, may also balance some losses in provinces with high levels of erosion (Schuerch et al., 2018).

A second hotspot for mangrove CO_2_ emissions was identified in the Tropical Northwest Atlantic, which had large emissions due to erosion (191 TgCO_2 eq_), clearing (80 TgCO_2 eq_), and extreme climatic events (23 TgCO_2 eq_), with total emissions projected to reach 312 TgCO_2 eq_ by the end of the century (Fig. 1, 2; Table S2). In the Mexican Caribbean, changes in hydrological connectivity that affect groundwater are a significant cause of unintended clearing of mangroves that are rich in carbon (Adame et al., 2013). The Tropical Northwest Atlantic is also one of the regions with the highest frequency of tropical storms in the world, which can cause large-scale mangrove mortality (Krauss & Osland, 2020).

Management activities to decrease CO_2_ emissions in the Tropical Northwest Atlantic could include the stabilisation of the coasts, reduction of illegal deforestation and the improvement of hydrological connectivity, especially in sites that fail to recover after tropical storms (Zaldívar-Jiménez et al., 2010). These activities could reduce the projected carbon emissions by 94% for this region (Fig. 3B).

Smaller hotspots with lower CO_2_ emissions were predicted to occur on the North Brazil Shelf, Sahul Shelf, and Gulf of Guinea (Fig. 1A), regions with an intermediate mangrove area and moderate carbon stocks (Table S1). In Brazil, vegetation clearing, changes in hydrology, and coastal development have increased erosion which has led to mangrove loss (Krause & Soares, 2004). Across the Sahul Shelf, northern Australia, the loss of mangroves during 2015-2016 was associated with an intense El Niño event which caused fluctuating sea levels, drought, and high temperatures (Lovelock, et al., 2017). In the Gulf of Guinea, CO_2_ emissions are the result of drought and changes in hydrology, which caused the loss of large areas of mangroves in Senegal (Sakho et al., 2017).

### Sensitivity of predictions to input data sources

The sensitivity analysis demonstrated that our projected hotspots of CO_2_ emissions due to mangrove loss are robust to different input data for mangrove area, carbon stock and emissions factors for extreme climatic events (Fig. S4). However, the total amount of emissions from SOC and ABC varied when we considered modelled versus field data sets, and different datasets for mangrove distribution (Fig S4). We considered field estimates to be more accurate, and since estimates to 2 m were closer to field SOC measurements than the estimates to 1 m (Fig. S2), we used 2 m as the depth of SOC for our predictions. Global emissions predictions based on the mangrove distribution dataset of Global Mangrove Watch (Bunting et al., 2018) were higher than those derived from the Hamilton and Casey (2016) dataset (Fig. S4). The former is considered a more comprehensive representation of mangrove forests globally because it captures mangroves of short stature. For instance, we found that predictions in provinces where short-statured mangroves are dominant (e.g. Tropical Northwest Atlantic) almost tripled when using the mangrove area from Global Mangrove Watch (Bunting et al., 2018).

The sensitivity analysis also indicated the model was most sensitive to the deforestation rates, with emissions increasing linearly as deforestation rate increased. The model was also sensitive to emission rates, but only in the short-term (Fig. S5-S8). Therefore, our model may overestimate emissions in regions where mangrove deforestation rates are slowing because of policy changes (Friess, et al., 2020; Richards et al., 2020). We further assumed that future rates of loss due to each of the five drivers were proportional to their historical contributions. Therefore, if deforestation rates continue to slow into the future, our CO_2_ emissions predictions will be overestimated. Changes in the magnitude of all land-use categories are likely to occur in the future, implying that our assumption of linearity in predictions may not happen. For example, unused agricultural land may transition to urban settlements. Also, mangrove loss may accelerate in some areas because of sea-level rise and climatic events. In other areas, the expansion of mangroves onto floodplains could compensate for some of the losses (Schuerch et al., 2018).

Overall our sensitivity analysis suggests that predictions on future emissions will be most sensitive to deforestation rates; thus, a research priority is developing scenarios for future mangrove loss that consider both climate and economic drivers of mangrove loss (Duarte et al., 2020; Schuerch et al., 2018). Our CO_2_ emission predictions could then be updated to account for changes in land-use trajectories – and the resulting changes in losses and gains – when higher-resolution global models of landscape change become available.

## Conclusion

We have identified hotspots of CO_2_ emissions due to mangrove loss associated with various drivers of land-use change. We predict that emissions arising from mangrove loss within this century will be concentrated in six provinces of the world: West Coral Triangle, Sunda Shelf, Bay of Bengal, Tropical Northwest Atlantic, Andaman, and North Brazil Shelf. These regions have large areas of mangroves (> 500,000 ha), relatively high rates of loss (≥ 0.1 % annually), and most of them have high carbon densities (≥ 500 MgC ha^−1^). By accounting for specific land-use changes and the foregone carbon sequestration potential, we update previous global estimates and provide specific management actions to most efficiently minimise future emissions. Activities that improve agricultural practices to reduce further expansion into mangrove areas, and efforts to stabilise coastlines and restore former mangrove areas should be prioritised to decrease emissions from mangrove loss by the end of the century.

## Acknowledgements

We acknowledge support from the Global Wetlands Project, supported by a charitable organisation which neither seeks nor permits publicity for its efforts. We thank the Global Mangrove Alliance and World Wildlife Foundation for guidance on this research project. CJB, RMC and MT are funded by a Discovery Project (DP180103124) from the Australian Research Council.

## Authors Contributions

MFA, RMC, CJB designed the project with major contributions from DF and CL; CJB, MT, and JH, designed the codes and models: LF, DL and LG, provided spatial data and analyses of mangrove loss drivers, JS provided spatial data and analyses of soil carbon, BK provided field data, SDF and DF provided data on emission factors; MS and DB conducted spatial analyses. MFA wrote the first draft and all authors contributed to editing subsequent drafts.

## Supplementary Material is available for this paper.

## Data availability

Data will be fully available at the Griffith University Data Repository and the repository of the Center for International Forestry Research (CIFOR).

