## Supplementary information for "Future carbon emissions from global mangrove forest loss"

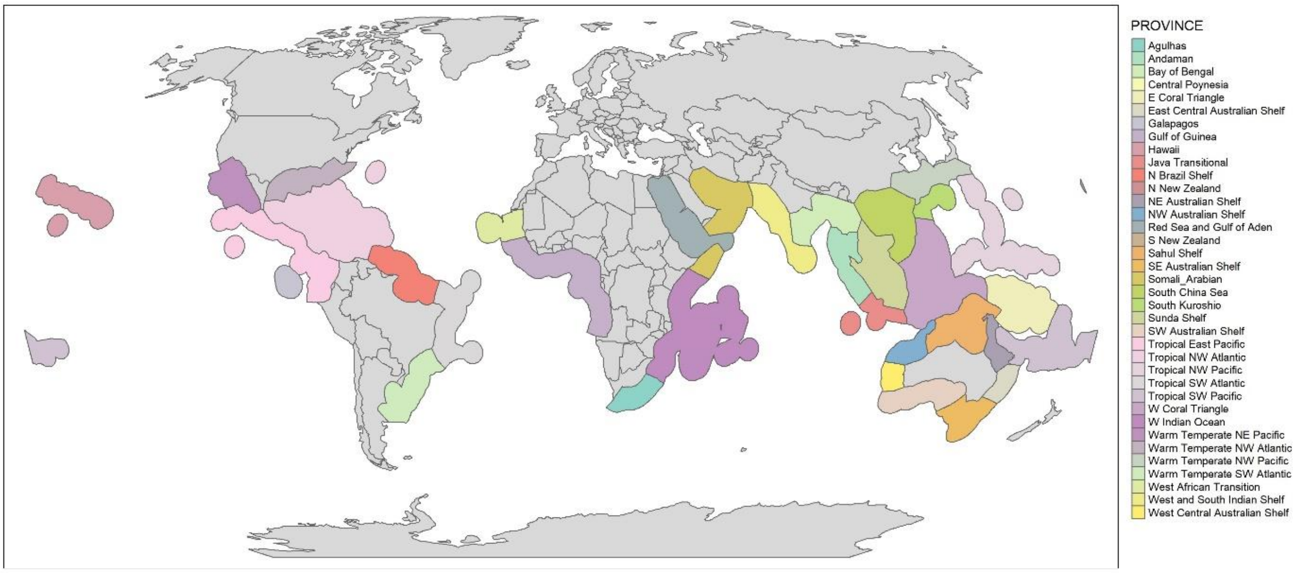


**Figure S1.** Marine provinces of the world with mangrove forests (Spalding et al., 2007).

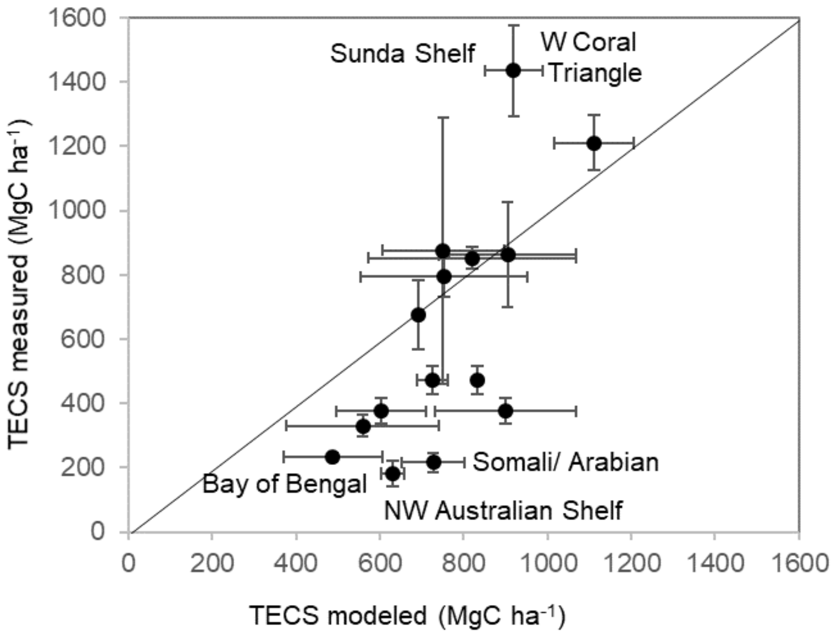


**Figure S2.** Relationship between modeled (ABC + SOC at 2m, mean ± SE among bioregions within one province) versus total ecosystem carbon stocks (TECS = ABC + SOC whole sediment column, mean ± SE among sites within one province) measured in the field (Kauffman et al., 2020) for 15 marine provinces of the world ($y$ = 1.90$x$ – 664, $R^2$ = 0.52, p = 0.002). Models slightly overestimate low TECS and underestimate large TECS. Continuous line indicates 1:1 relationship

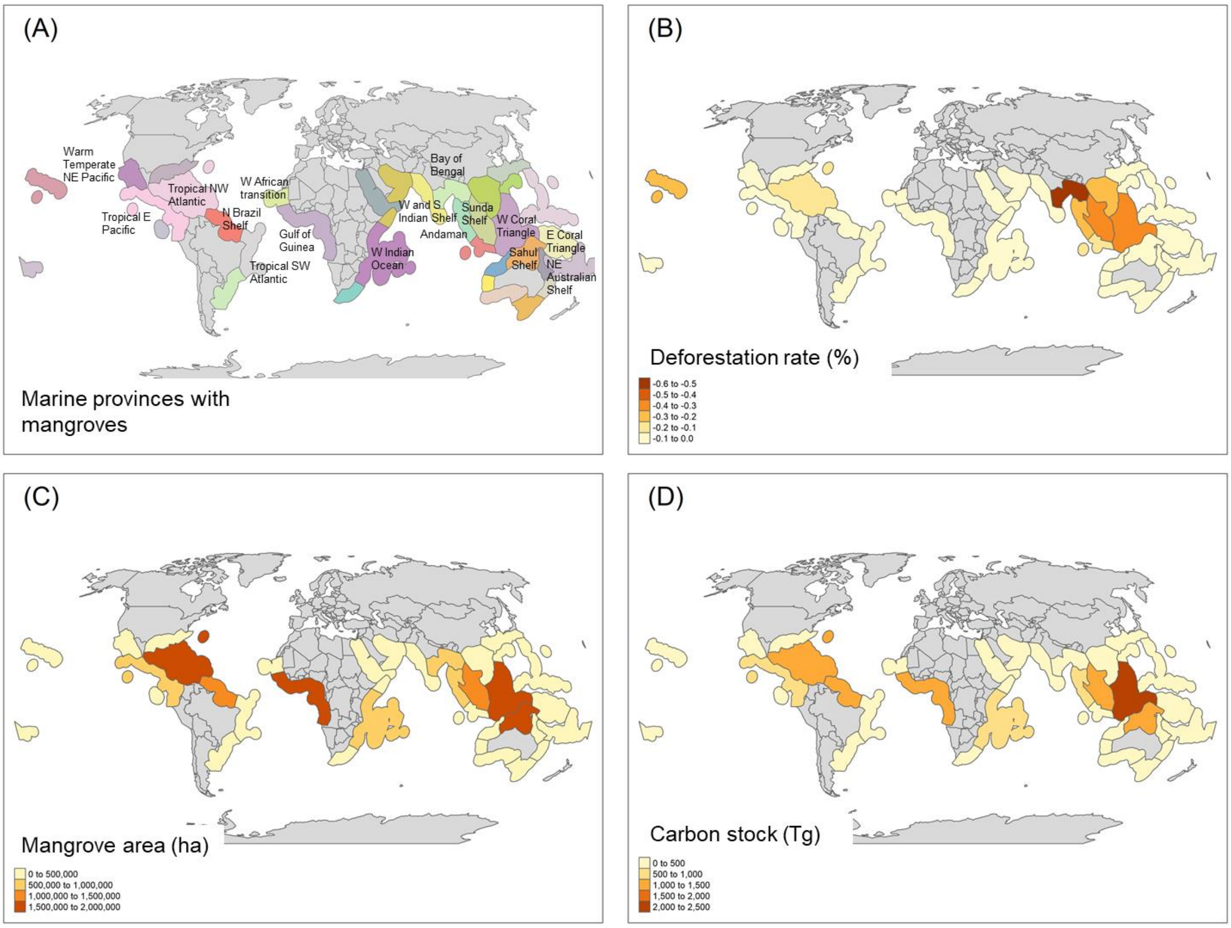


**Figure S3**. (A) Marine provinces of the world with mangroves (see Table S1 for a list of all provinces and Fig S1. for a map with the labels of all provinces), B) deforestation rates (2000-2012, %) (Hamilton & Casey, 2016), (C) mangrove area in 2010 (ha) (Bunting et al., 2018), and (D) total carbon stored (Tg) (Sanderman et al., 2018; Simard et al. 2019).

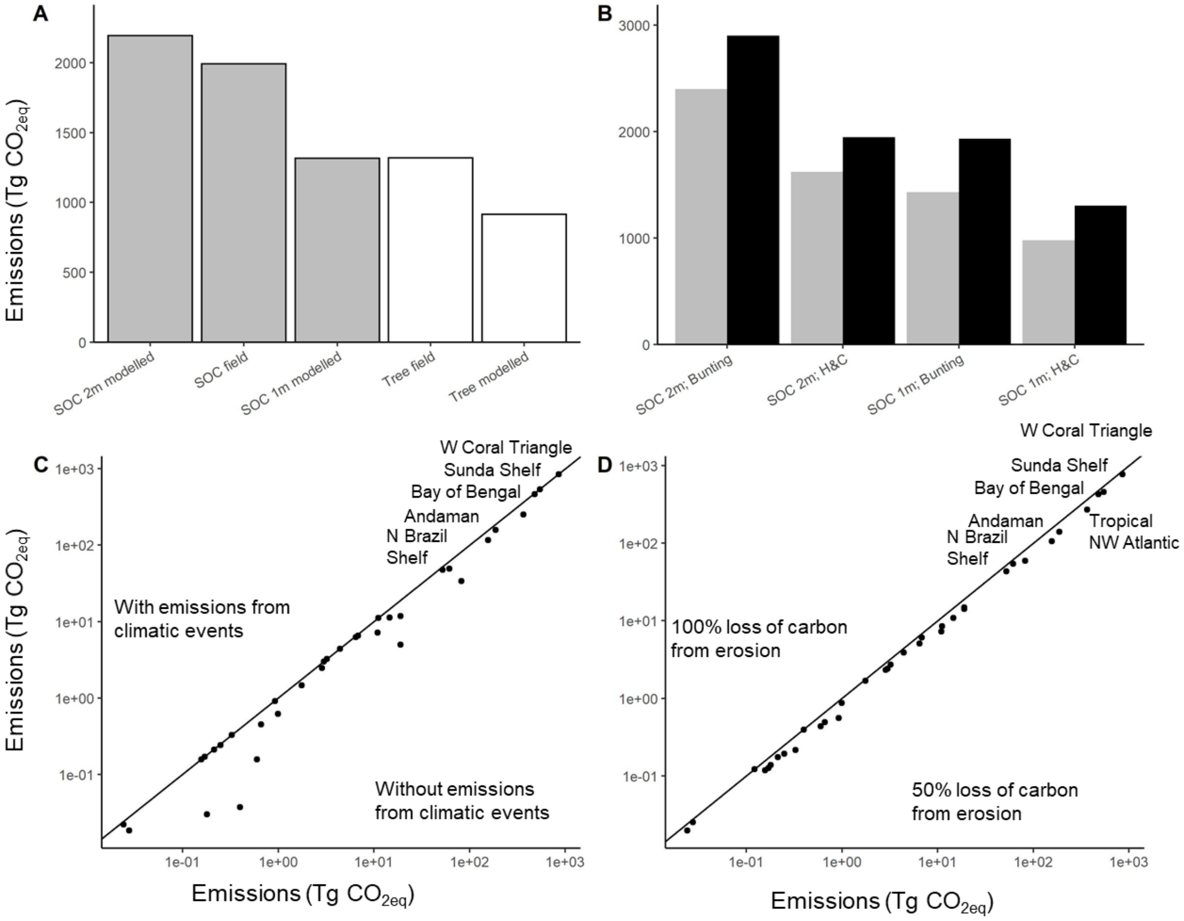


**Figure S4.** Sensitivity analyses comparing cumulative $CO_{2eq}$ emissions at the end of the century (2010-2100) using A) soil organic carbon (SOC) modeled (1 and 2 m deep) (Sanderman et al., 2018), SOC obtained in the field (Kauffman et al., 2020), and modeled aboveground biomass carbon (Tree) (Simard et al., 2019) and obtained in the field for the 15 provinces with field data (Kauffman et al., 2020); B) comparison between emissions with mangrove area from Bunting et al (2018) and with Hamilton and Casey (H&C) (2016) with SOC at 1 and 2m; (C) comparison of ranking among provinces with highest cumulative $CO_{2eq}$ emissions with and without accounting for emissions from climate events, and (D) with low (0.5) and high (1) emission factors from erosion.

**Table S1.** Mangrove area baseline (2010) (Bunting et al., 2018) for marine provinces (Spalding et al., 2007), annual deforestation rate (DR %, 2000-2012) (Hamilton & Casey, 2016), soil organic carbon stocks (SOC, MgC $ha^{-1}$) for the top one and two meters of soil (Sanderman et al., 2018), aboveground biomass carbon stocks (ABC, MgC $ha^{-1}$) (Simard et al., 2019), and total ecosystems carbon stocks from modeled data (TECS, MgC $ha^{-1}$, ABC + SOC 2m) and from direct measurements in the field (TECS-field, MgC $ha^{-1}$) (Kauffman et al., 2020).

| Marine Province | Area (ha) | DR (%) | SOC 1m | SOC 2m | ABC | TECS | TECS-field |
|---|---|---|---|---|---|---|---|
| | | | (MgC $ha^{-1}$) | | | | |
| West Coral Triangle | 1,836,289 | 0.33 | 497.8 | 975.9 | 134.3 | 1110.2 | 1210.0 |
| Gulf of Guinea | 1,806,989 | 0.02 | 341.4 | 665.2 | 87.7 | 752.9 | 794.0 |
| Sahul Shelf | 1,656,126 | 0.03 | 399.8 | 794.5 | 103.3 | 897.8 | 376.0 |
| Tropical Northwest Atlantic | 1,534,395 | 0.14 | 332.4 | 827.3 | 78.2 | 905.5 | 863.4 |
| North Brazil Shelf | 1,263,556 | 0.06 | 362.7 | 724.4 | 107.1 | 831.5 | 473.0 |
| Sunda Shelf | 1,127,741 | 0.35 | 408.0 | 815.4 | 103.2 | 918.6 | 1434.9 |
| Bay of Bengal | 911,223 | 0.55 | 207.4 | 408.6 | 78.8 | 487.4 | 234.0 |
| West Indian Ocean | 728,044 | 0.02 | 335.7 | 657.6 | 92.7 | 750.3 | 875.7 |
| Tropical East Pacific | 727,814 | 0.04 | 416.4 | 650.1 | 169.7 | 819.8 | 852.8 |
| Andaman | 500,856 | 0.22 | 466.9 | 911.8 | 100.1 | 1011.9 | |
| Northeast Australian Shelf | 232,406 | 0.03 | 292.0 | 564.9 | 44.7 | 609.6 | 376.0 |
| Warm Temperate Northeast Pacific | 223,721 | 0.02 | 262.6 | 503.6 | 55.8 | 559.4 | 331.0 |
| East Coral Triangle | 215,918 | 0.06 | 468.9 | 919.6 | 109.1 | 1028.7 | |
| West African Transition | 184,566 | 0.002 | 318.4 | 622.8 | 67.4 | 690.2 | 675.0 |
| Tropical Southwest Atlantic | 171,645 | 0.03 | 328.2 | 638.9 | 86.0 | 724.9 | 473.0 |
| West and South Indian Shelf | 143,134 | 0.07 | 284.2 | 558.8 | 61.5 | 620.3 | |
| Tropical Southwest Pacific | 85,480 | 0.02 | 406.1 | 790.4 | 79.8 | 870.2 | |
| Warm Temperate Southwest Atlantic | 73,236 | 0.01 | 403.9 | 781.6 | 113.2 | 894.8 | |
| Northwest Australian Shelf | 47,831 | 0.01 | 316.0 | 615.3 | 15.4 | 630.7 | 181.5 |
| East Central Australian Shelf | 40,550 | 0.04 | 356.6 | 502 | 108.5 | 610.5 | |
| South China Sea | 39,706 | 0.22 | 237.8 | 547.0 | 52.7 | 509.7 | |
| Southwest Australian Shelf | 19,883 | 0.01 | 284.1 | 539.4 | 9.8 | 549.2 | |
| Somali/Arabian | 18,852 | 0.001 | 252.7 | 495.2 | 220.7 | 715.9 | 217.0 |

| | | | | | | |
|---|---|---|---|---|---|---|
| North New Zealand | 16,063 | 0.10 | 427.8 | 823.3 | 35.3 | 858.6 |
| Java Transitional | 15,119 | 0.17 | 483.2 | 954.6 | 131.8 | 1086.4 |
| Red Sea and Gulf of Aden | 15,066 | 0.03 | 272.8 | 538.1 | 199.0 | 737.1 |
| South New Zealand | 14,534 | 0.10 | 427.8 | 823.3 | 103.8 | 927.1 |
| Tropical Northwest Pacific | 14,490 | 0.03 | 578.9 | 1155.8 | 56.5 | 1212.3 |
| Southeast Australian Shelf | 5,022 | 0.03 | 264.4 | 516.9 | 20.3 | 537.2 |
| Warm Temperate Northwest Atlantic | 3,728 | 0.04 | 404.7 | 808.1 | 61.9 | 870.0 |
| Agulhas | 2,581 | 0.09 | 239.2 | 457.9 | 86.0 | 543.9 |
| Galapagos | 2,450 | 0.005 | 426.9 | 829.6 | 169.7 | 999.3 |
| West Central Australian Shelf | 1,671 | 0 | 277.5 | 529.5 | 25.5 | 555.0 |
| Warm Temperate Northwest Pacific | 1,228 | 0.08 | 291.4 | 561.0 | 63.2 | 624.2 |
| South Kuroshio | 983 | 0.06 | 411.8 | 801.1 | 332.3 | 1133.4 |
| Hawaii | 704 | 0.27 | 275.4 | 532.4 | 80.8 | 613.2 |
| Central Polynesia | 324 | 0 | 406.1 | 790.4 | 44.2 | 834.6 |

**Table S2.** Emission factors (Sasmito et al., 2019) for aboveground biomass carbon (ABC), soil organic carbon (SOC), and level of confidence (Tier), where: Level 1 is a value obtained from the global average, Level 2 is a value from a similar region, and Level 3 is a value from a similar region and geomorphic setting.

| Marine Province | Agriculture/aquaculture | | | | Erosion | | | Clearing | | | | Extreme Climate | | | Settlement | | |
|---|---|---|---|---|---|---|---|---|---|---|---|---|---|---|---|---|---|
| | ABG | Tier | SOC | Tier | ABG | SOC | Tier | ABG | Tier | SOC | Tier | ABG | SOC | Tier | ABG | SOC | Tier |
| Agulhas | 0.83 | 1 | 0.52 | 1 | 1 | 1 | 1 | 0.70 | 1 | 0.21 | 3 | 0.31 | 0.14 | 1 | 1 | 0.66 | 1 |
| Andaman | 0.90 | 2 | 0.27 | 3 | 1 | 1 | 1 | 0.88 | 3 | 0.45 | 3 | 0.31 | 0.14 | 1 | 1 | 0.66 | 1 |
| Bay of Bengal | 0.83 | 1 | 0.52 | 1 | 1 | 1 | 1 | 0.70 | 1 | 0.33 | 1 | 0.31 | 0.14 | 1 | 1 | 0.66 | 1 |
| East Central Australian Shelf | 0.83 | 1 | 0.52 | 1 | 1 | 1 | 1 | 0.70 | 1 | 0.33 | 1 | 0.31 | 0.14 | 1 | 1 | 0.66 | 1 |
| East Coral Triangle | 0.83 | 1 | 0.52 | 1 | 1 | 1 | 1 | 0.70 | 1 | 0.33 | 1 | 0.31 | 0.14 | 1 | 1 | 0.66 | 1 |
| Gulf of Guinea | 0.83 | 1 | 0.52 | 1 | 1 | 1 | 1 | 0.70 | 1 | 0.33 | 1 | 0.31 | 0.14 | 1 | 1 | 0.66 | 1 |
| North Brazil Shelf | 0.97 | 2 | 0.67 | 3 | 1 | 1 | 1 | 0.70 | 1 | 0.33 | 1 | 0.31 | 0.14 | 2 | 1 | 0.66 | 1 |
| Northeast Australian Shelf | 0.83 | 1 | 0.52 | 1 | 1 | 1 | 1 | 0.70 | 1 | 0.33 | 1 | 0.31 | 0.14 | 1 | 1 | 0.66 | 1 |
| North New Zealand | 0.83 | 1 | 0.52 | 1 | 1 | 1 | 1 | 1.00 | 3 | 0.33 | 1 | 0.31 | 0.14 | 1 | 1 | 0.66 | 1 |
| Northwest Australian Shelf | 0.83 | 1 | 0.52 | 1 | 1 | 1 | 1 | 0.70 | 1 | 0.33 | 1 | 0.31 | 0.14 | 1 | 1 | 0.66 | 1 |
| Red Sea and Gulf of Aden | 0.83 | 1 | 0.52 | 1 | 1 | 1 | 1 | 0.70 | 1 | 0.33 | 1 | 0.31 | 0.14 | 1 | 1 | 0.66 | 1 |
| Sahul Shelf | 0.90 | 2 | 0.27 | 3 | 1 | 1 | 1 | 0.88 | 3 | 0.45 | 3 | 0.31 | 0.14 | 1 | 1 | 0.66 | 1 |
| Somali/Arabian | 0.83 | 1 | 0.52 | 1 | 1 | 1 | 1 | 0.88 | 2 | 0.45 | 2 | 0.31 | 0.14 | 1 | 1 | 0.66 | 1 |
| South China Sea | 0.83 | 1 | 0.52 | 1 | 1 | 1 | 1 | 0.88 | 3 | 0.45 | 3 | 0.31 | 0.14 | 2 | 1 | 0.66 | 1 |
| Southeast Australian Shelf | 0.83 | 1 | 0.52 | 1 | 1 | 1 | 1 | 0.88 | 3 | 0.45 | 3 | 0.31 | 0.14 | 1 | 1 | 0.66 | 1 |

| | | | | | | | | | | | | | | | | | |
|---|---|---|---|---|---|---|---|---|---|---|---|---|---|---|---|---|---|
| Southern New Zealand | 0.83 | 1 | 0.52 | 1 | 1 | 1 | 1 | 1.00 | 2 | 0.60 | 3 | 0.31 | 0.14 | 1 | 1 | 0.66 | 1 |
| Southwest Australian Shelf | 0.83 | 1 | 0.52 | 1 | 1 | 1 | 1 | 0.88 | 2 | 0.45 | 2 | 0.31 | 0.14 | 1 | 1 | 0.66 | 1 |
| Sunda Shelf | 0.90 | 2 | 0.27 | 3 | 1 | 1 | 1 | 0.88 | 3 | 0.45 | 3 | 0.31 | 0.14 | 1 | 1 | 0.66 | 1 |
| Tropical East Pacific | 0.83 | 1 | 0.52 | 1 | 1 | 1 | 1 | 0.88 | 2 | 0.45 | 3 | 0.31 | 0.14 | 1 | 1 | 0.66 | 1 |
| Tropical Northwestern Atlantic | 0.76 | 2 | 0.46 | 3 | 1 | 1 | 1 | 0.88 | 2 | 0.45 | 2 | 0.31 | 0.14 | 1 | 1 | 0.66 | 1 |
| Warm Temperate Northeast Pacific | 0.83 | 1 | 0.52 | 1 | 1 | 1 | 1 | 0.88 | 2 | 0.45 | 2 | 0.31 | 0.14 | 1 | 1 | 0.66 | 1 |
| Warm Temperate Southwestern Atlantic | 0.97 | 2 | 0.67 | 3 | 1 | 1 | 1 | 0.88 | 3 | 0.45 | 3 | 0.31 | 0.14 | 1 | 1 | 0.66 | 1 |
| West African Transition | 0.83 | 1 | 0.52 | 1 | 1 | 1 | 1 | 0.88 | 2 | 0.45 | 2 | 0.31 | 0.14 | 1 | 1 | 0.66 | 1 |
| West and South Indian Shelf | 1.00 | 2 | 0.45 | 3 | 1 | 1 | 1 | 0.88 | 2 | 0.45 | 2 | 0.31 | 0.14 | 1 | 1 | 0.66 | 1 |
| West Central Australian Shelf | 0.83 | 1 | 0.52 | 1 | 1 | 1 | 1 | 0.88 | 2 | 0.45 | 2 | 0.31 | 0.14 | 1 | 1 | 0.66 | 1 |
| Western Coral Triangle | 0.90 | 3 | 0.27 | 2 | 1 | 1 | 1 | 0.88 | 3 | 0.45 | 3 | 0.31 | 0.14 | 1 | 1 | 0.66 | 1 |
| Western Indian Ocean | 0.83 | 1 | 0.52 | 1 | 1 | 1 | 1 | 0.70 | 3 | 0.21 | 3 | 0.31 | 0.14 | 1 | 1 | 0.66 | 1 |

**Table S3.** Cumulative emissions ($TgCO_{2\ eq}$) projected for the next century (2020-2100) derived from drivers of land-use change: commodities (agriculture/aquaculture), erosion, clearing or non-productive conversion, extreme climatic events, and human settlements (Goldberg et al., 2020). The first six provinces account for 90% of global emissions.

| Marine Province | Agriculture/ aquaculture | Erosion | Clearing | Extreme climatic events | Human Settlement | Total |
|---|---|---|---|---|---|---|
| West Coral Triangle | 519.9 | 163.8 | 18.2 | 1.7 | 8.5 | 712.1 |
| Sunda Shelf | 221.3 | 173.4 | 14.7 | 1.8 | 40.6 | 451.8 |
| Bay of Bengal | 243.7 | 112.2 | 8.4 | 3.1 | 1.2 | 368.6 |
| Tropical Northwest Atlantic | 9.1 | 190.8 | 79.9 | 22.7 | 9.6 | 312.1 |
| Andaman | 41.9 | 97.5 | 14.4 | 6.4 | 1.2 | 161.4 |
| North Brazil Shelf | 21.9 | 103.5 | 4.1 | 7 | 0.1 | 136.6 |
| Sahul Shelf | 0.1 | 47.6 | 5.1 | 19 | 0 | 71.8 |
| Gulf of Guinea | 5.7 | 14.6 | 15.5 | 2.5 | 11.3 | 49.6 |
| Tropical E Pacific | 18.7 | 17.7 | 5.3 | 0.8 | 2.3 | 44.8 |
| East Coral Triangle | 0.1 | 9.8 | 1.3 | 4.3 | 0.1 | 15.6 |
| West Indian Ocean | 1.3 | 8.2 | 3.3 | 1.7 | 0.3 | 14.8 |
| West and S Indian Shelf | 0.7 | 7.4 | 2.4 | 0.6 | 0.8 | 11.9 |
| South China Sea | 2.4 | 5.3 | 0.5 | 0 | 0.8 | 9 |
| Northeast Australian Shelf | 0 | 7.5 | 0.8 | 0.7 | 0 | 9 |
| Warm Temperate Southwest Atlantic | 3.4 | 2.4 | 0.5 | 0.8 | 1.3 | 8.4 |
| Tropical Southwest Atlantic | 2.4 | 1.5 | 1.5 | 0 | 0.2 | 5.6 |
| Warm Temperate Norhteast Pacific | 1.5 | 2.8 | 0.8 | 0 | 0 | 5.1 |
| Java Transitional | 1.5 | 1.1 | 0.8 | 0 | 0.4 | 3.8 |
| Tropical Southwesta Pacific | 0 | 1 | 1.2 | 0.1 | 0 | 2.3 |
| East Central Australian Shelf | 0 | 0.2 | 0.9 | 0.1 | 0.2 | 1.4 |
| West African Transition | 0 | 0.7 | 0.1 | 0 | 0 | 0.8 |
| Red Sea and Gulf of Aden | 0.1 | 0.3 | 0.1 | 0 | 0 | 0.5 |
| Tropical Northwest Pacific | 0 | 0.3 | 0 | 0.1 | 0 | 0.4 |
| Northwest Australian Shelf | 0 | 0 | 0.1 | 0.2 | 0 | 0.3 |
| Agulhas | 0 | 0.2 | 0 | 0 | 0 | 0.2 |
| Hawaii | 0.1 | 0.1 | 0 | 0 | 0 | 0.2 |
| Southwest Australian Shelf | 0 | 0.1 | 0.1 | 0 | 0 | 0.2 |
| South Kuroshio | 0 | 0.1 | 0 | 0 | 0 | 0.1 |
| Warm Temperate NW Pacific | 0 | 0.1 | 0 | 0 | 0 | 0.1 |
| Southeast Australian Shelf | 0 | 0 | 0 | 0.1 | 0 | 0.1 |
| Warm Temperate Northwest Atlantic | 0 | 0 | 0 | 0 | 0 | 0 |
| Somali/Arabian | 0 | 0 | 0 | 0 | 0 | 0 |

| | | | | | | |
|---|---|---|---|---|---|---|
| Galapagos | 0 | 0 | 0 | 0 | 0 | 0 |
| West Central Australian Shelf | 0 | 0 | 0 | 0 | 0 | 0 |
| TOTAL | 1,095.8 | 970.2 | 180.0 | 78.9 | 73.7 | 2,398.6 |

### Supplementary Text

Model derivation

The model sensitivity analyses were plotted for the median, 1st quartile and 3rd quartile of the data (Table S4

**Table S4.** The median, 1st quartile and 3rd quartile of the data used to parameterize the model.

| | Median | 1st Quartile | 3rd Quartile |
|---|---|---|---|
| Initial area $A_1$ (Ha) | 178 603 | 23 047 | 875 276 |
| Deforestation rate d (per year) | 0.00193 | 0.00115 | 0.00265 |
| Carbon storage/maximum emissions per hectare c (Mg $C0_2$/Ha) | 923 | 749 | 1091 |
| Emission rate r (per year) | 0.1 | 0.1 | 0.1 |
| Sequestration rate s (Mg $C0_2$ /year) | 6.49 | 6.49 | 6.49 |

We assumed mangroves in a marine province were lost at a constant rate:

$$\frac{dA}{dt} = -Ad \tag{1}$$

Where $A$ is the Area at time ($t$) and $d$ is the rate of loss. The area of mangroves lost ($D_t$) is equal to the difference in initial Area ($A_1$) and Area at a specific time ($A_t$):

$$D_t = A_1(1 - e^{-dt}) \tag{2}$$

From the above, we derived the rate of change in the area of mangrove lost:

$$\frac{dD_t}{dt} = A_1 \cdot d \cdot e^{-dt} \tag{3}$$

The $CO_2$ emissions from mangrove loss depend on the area and the rate of emissions. Therefore, we considered two separate time quantities: $t$ is the total time since loss began anywhere in the marine province and *y* is the instant when a specific area lost its mangroves. Hence the number of years of emissions from an area of deforested mangrove is *t*-*y*.

The rate of change in $CO_2$ emissions with respect to year mangrove loss began is therefore dependent on the rate of change in the area and the rate of emissions ($r$) up to a limit for maximum carbon that could be emitted per hectare of forest ($c$).

$$\frac{dE}{dy} = A_1 \cdot d \cdot e^{-dt} \cdot c \cdot r \cdot e^{-(t-y)r} \tag{4}$$

Emissions rates were linked to the length of time since mangrove loss occurred, so emissions were scaled by: $e^{-(t-y)r}$

To find emissions at year *t*, we integrate over *y*:

$$E = \int_0^y A_1 \cdot d \cdot e^{-dt} \cdot c \cdot r \cdot e^{-(t-y)r} dy$$

$$E_t = \frac{A_1 \cdot d \cdot c \cdot r \cdot e^{-tr} \cdot \left(e^{(r-d)t} - 1\right)}{(r-d)} \quad \text{for r≠d} \tag{5}$$

The above model predicted that $CO_2$ emissions were undefined when $r = d$, because the denominator was equal to 0. We solved for emissions when $r=d$ by applying l'hopitals rule to find emissions in the limit of $r$ approaching $d$:

$$E_t = \lim_{r \to d} \frac{A_1 \cdot d \cdot c \cdot r \cdot e^{-tr} \cdot (e^{(r-d)t} - 1)}{(r-d)}$$

$$E_t = A_1 \cdot d^2 \cdot c \cdot t \cdot e^{-td} \quad \text{for r=d} \tag{6}$$

Finally, we integrated both equations 5 and 6 over time to calculate the cumulative emissions ($J_t$) from deforestation at a year, giving:

$$J_t = \int_0^t \frac{A_1 \cdot d \cdot c \cdot r \cdot e^{-tr} \cdot \left(e^{(r-d)t} - 1\right)}{(r-d)} dt$$

$$J_t = \frac{A_1 \cdot c \cdot (d \cdot e^{-rt} - r \cdot e^{-dt} + (r-d))}{(r-d)} \text{ if r} \neq \text{d} \tag{7}$$

$$J_t = A_1 \cdot c \cdot (1 - e^{-dt} - d \cdot t \cdot e^{-dt}) \text{ if r=d} \tag{8}$$

An example of the $CO_2$ emissions (instant and cumulative) is shown in Figure S5.

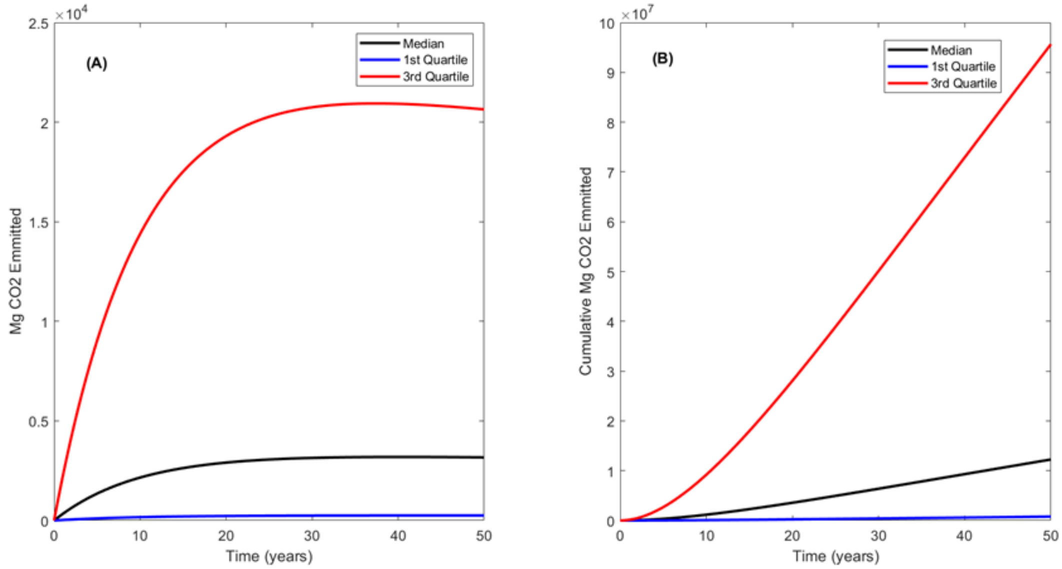


**Figure S5.** (A) Emissions (Mg $CO_{2eq}$, median 1st and 3rd quartile) at a point in time, and (B) cumulative emission over time

Carbon sequestration

Sequestered carbon (*S*) was assumed to have a constant rate of accumulation per hectare of mangrove forest (Adame et al., 2018):

$$\frac{dS}{dt} = s \cdot A_1 e^{-dt} \tag{9}$$

Where *s* was the sequestration rate (Mg ha$^{-1}$) and *S* was the component of carbon that had been sequestered since the initial year. Therefore:

$$S_t = \int s \cdot A_1 e^{-dt} dt$$

$$S_t = \frac{-A_1 s e^{-dt}}{d} + constant \tag{10}$$

At $t = 0$ the carbon is equal to $C_1$ (initial carbon within the soil), therefore the constant can be solved to equal $C_1 + \frac{A_1 s}{d}$.

The $C_1$ can be ignored, since we are only interested in sequestration since year 0:

$$S_t = \frac{A_1 s\,(1 - e^{-dt})}{d} \quad \text{for } d \neq 0$$

(11)

$$S_t = A_1 st \text{ for d=0}$$

(12)

Total carbon stored ($C_t$) in the mangroves at a time is the sum of initial storage ($C_1$), sequestration ($S_t$) and emissions losses ($J_t$):

$$C_t = S_t + C_1 - J_t$$

$$C_t = \frac{A_1 s\,(1 - e^{-dt})}{d} + C_1 - \frac{A_1 \cdot c \cdot \left(d \cdot e^{-rt} - r \cdot e^{-dt} + (r-d)\right)}{(r-d)} \quad \text{if r} \neq \text{d and d} \neq 0$$

(13)

$$C_t = \frac{A_1 s\,(1 - e^{-dt})}{d} + C_1 - A_1 \cdot c \cdot (1 - e^{-dt} - d \cdot t \cdot e^{-dt}) \quad \text{if r = d and d} \neq 0$$

(14)

A complete account of $CO_2$ emissions and sequestration would include emissions from mangrove loss and the forgone opportunity to sequester carbon caused by mangrove loss. We therefore calculate the carbon account as the difference between carbon storage with mangrove loss and a counterfactual for no loss:

$$L_t = C_t^0 - C_t^d$$

$$L_t = S_t^0 - S_t^d + J_t^d$$

$$L_t = A_1 st - \frac{A_1 s\,(1 - e^{-dt})}{d} + \frac{A_1 c(d\, e^{-rt} - r\, e^{-dt} + (r-d))}{(r-d)} \quad \text{for r} \neq \text{d}$$

(15)

$$L_t = A_1 st - \frac{A_1 s\,(1 - e^{-dt})}{d} + A_1 \cdot c \cdot (1 - e^{-dt} - d \cdot t \cdot e^{-dt}) \quad \text{for r=d}$$

(16)

Where $C_t^d$ is carbon storage with mangrove loss and $C_t^0$ is carbon storage with zero net loss. When $s$=0 equations 13 and 14 revert to the negative of the emissions equation ($J_t$).

Sensitivity of the model

We studied the sensitivity of Equation 15 to timescale (50 years) and its parameters by finding the derivatives of the equation with respect to each parameter. We also studied the special case of $s = 0$, which reflects a carbon account where we only include emissions and not lost opportunity to sequester carbon. The sensitivity of the lost opportunity to store carbon to each parameter was:

$$L_t = -\frac{A_1 s\,(1 - e^{-dt})}{d} + \frac{A_1 c\left(d\,e^{-rt} - r\,e^{-dt} + (r-d)\right)}{(r-d)} + A_1 st \text{ for } r \neq d \tag{17}$$

$$\frac{dL_t}{ds} = -\frac{A_1(1 - e^{-dt})}{d} + A_1 t \text{ for } d \neq 0 \tag{18}$$

$$\frac{dL_t}{dc} = \frac{A_1(de^{-rt} - re^{-dt} + r - d)}{(r-d)} \text{ for } r \neq d \tag{19}$$

$$\frac{dL_t}{dd} = \frac{A_1 c(de^{-rt} - re^{-dt} + r - d)}{(r-d)^2} + \frac{A_1 c(e^{-rt} - rte^{-dt} - 1)}{(r-d)} + \frac{A_1 s(1 - e^{-dt})}{d^2} - \frac{A_1 ste^{-dt}}{d} \text{ for } r \neq d \tag{20}$$

$$\frac{dL_t}{dr} = -\frac{A_1 c(de^{-rt} - re^{-dt} + r - d)}{(r-d)^2} + \frac{A_1 c(-dte^{-rt} - e^{-dt} + 1)}{(r-d)} \text{ for } r \neq d \tag{21}$$

The rate of soil sequestration and the maximum carbon storage increased in importance at longer time scales (Fig. S6A, B), but overall the model was not highly sensitive to these parameters. $L_t$ was most sensitive to the rate of deforestation, as evidenced by the largest changes in $L_t$ on the y-axis compared to other parameters, increasing linearly with time (Fig. S6C). The rate of emissions was also important, but its importance decreased with time (Fig. S6D).

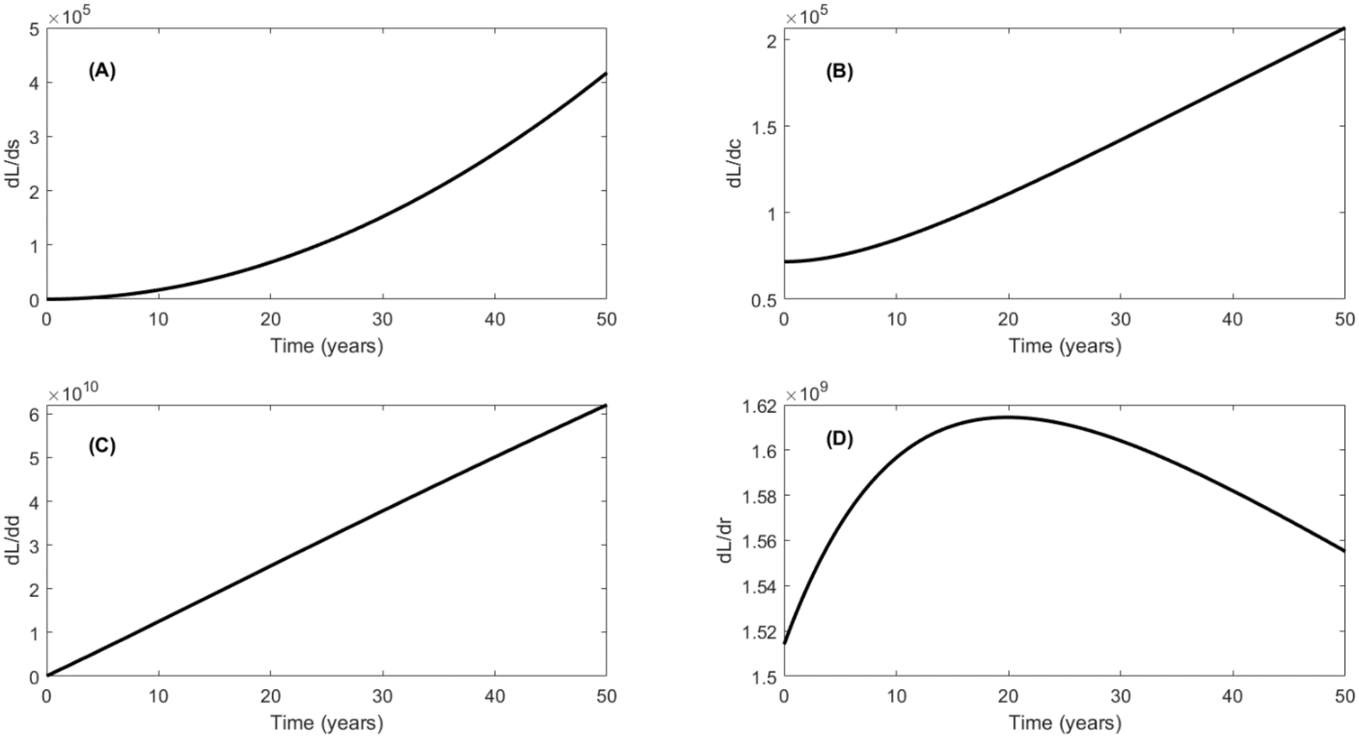


**Figure S6.** Sensitivity of model to changes in (A) soil sequestration rates, (B) maximum carbon loss, (C) deforestation rate, and (D) emission rate.

The sensitivity of the model on forgone soil carbon sequestration was compared with a model without carbon sequestration:

$$L_{t\,(s=0)} = \frac{A_1 c(d\,e^{-rt} - r\,e^{-dt} + (r-d))}{(r-d)} \text{ for } r \neq d \tag{22}$$

$$\frac{dL_{t\,(s=0)}}{dc} = \frac{A_1(de^{-rt} - re^{-dt} + r - d)}{(r-d)} \text{ for } r \neq d \tag{23}$$

$$\frac{dL_{t\,(s=0)}}{dd} = \frac{A_1 c(de^{-rt} - re^{-dt} + r - d)}{(r-d)^2} + \frac{A_1 c(e^{-rt} - rte^{-dt} - 1)}{(r-d)} \text{ for } r \neq d \tag{24}$$

$$\frac{dL_{t\,(s=0)}}{dr} = \frac{-A_1 c(de^{-rt} - re^{-dt} + r - d)}{(r-d)^2} + \frac{A_1 c(-dte^{-rt} - e^{-dt} + 1)}{(r-d)} \text{ for } r \neq d \tag{25}$$

Both models had similar sensitivities to maximum carbon storage, deforestation and emission rates (Fig. S6, S7).

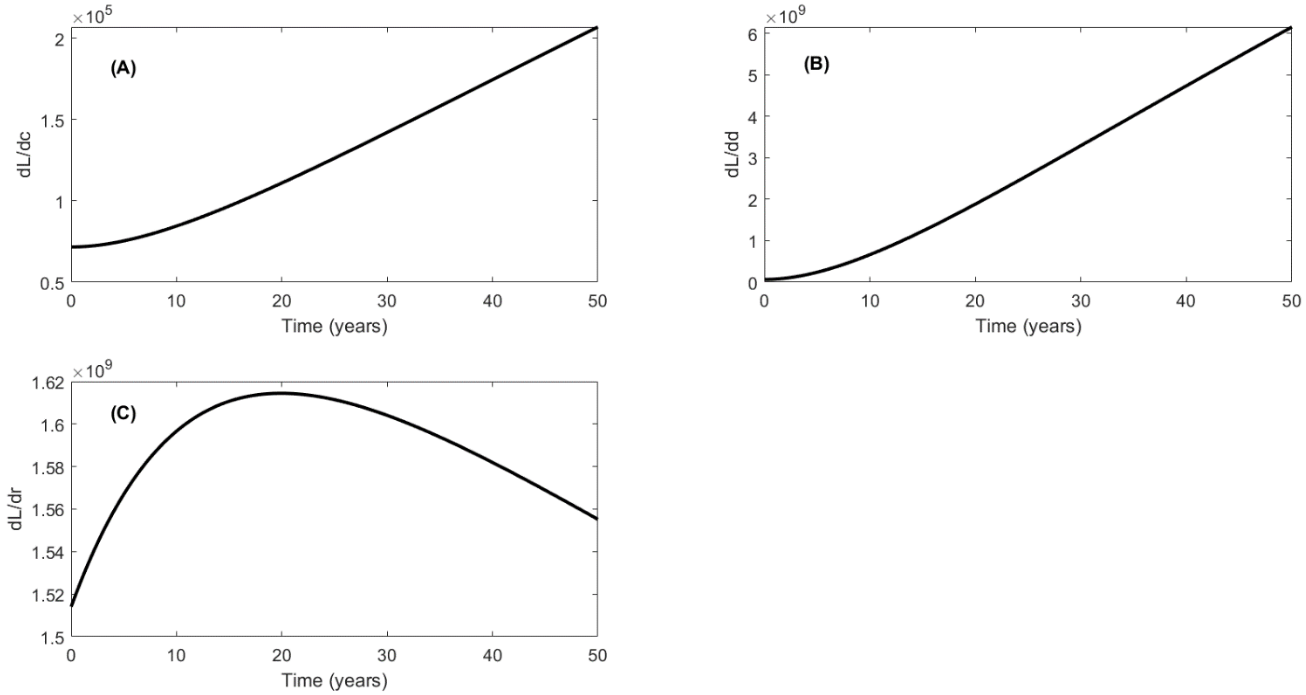


**Figure S7**. Sensitivity of model to changes in (A) maximum carbon loss, (B) deforestation, and (C) emission rate

Numerical sensitivity analysis

The parameters were varied from the lower quartile to the upper quartile to assess the sensitivity of forgone soil carbon to their variation (Table S4). The forgone soil carbon at 50 years increased close to linearly for higher sequestration, deforestation rates and higher maximum emissions value. The model was not sensitive to the emission rate at the timescale of 50 years (Fig. S8).

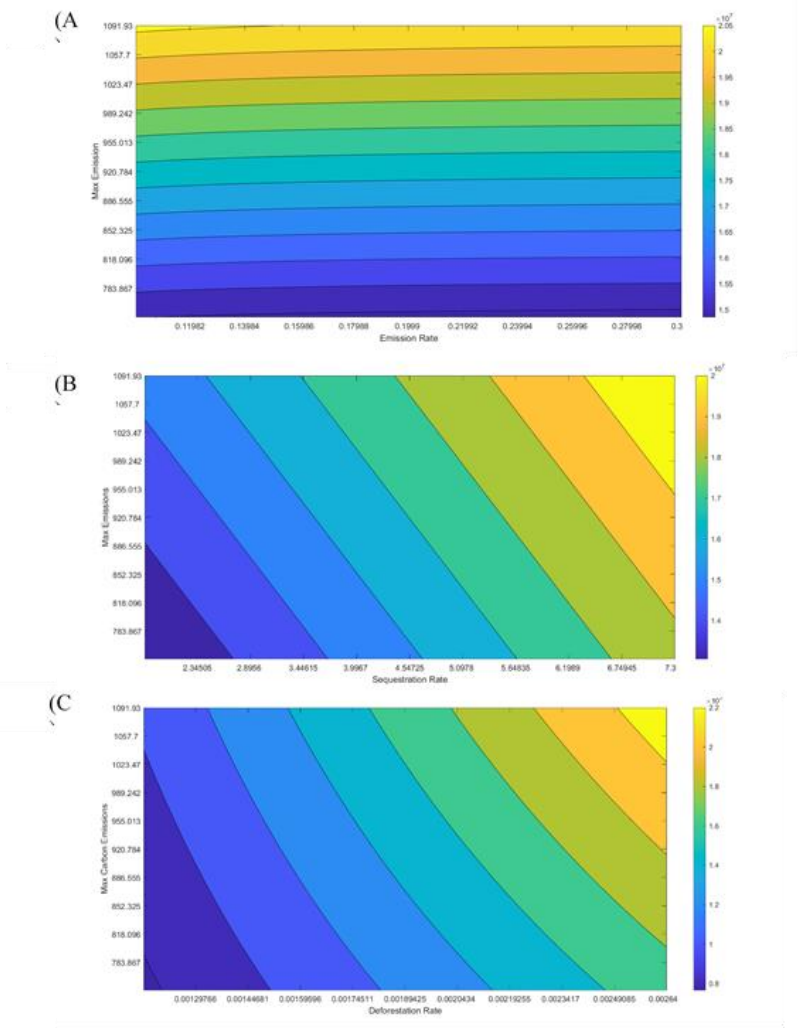


**Figure S8.** Sensitivity of the forgone soil carbon to (A) maximum emissions and the emission rate, (B) maximum emissions and the sequestration rate and (C) maximum emissions and the deforestation rate.
